# Three-dimensional T_1_ mapping demonstrates the transfer of oxygen into cerebrospinal fluid during hyperoxia

**DOI:** 10.64898/2025.12.01.691570

**Authors:** Emma Biondetti, Davide Di Censo, Sara Pomante, Stefano Censi, Ekaterina Bliakharskaia, Manuela Carriero, Lucie Chalet, Giulia Rocco, Guido Buonincontri, Francesca Graziano, Alessandra Stella Caporale, Antonio Maria Chiarelli, Richard Geoffrey Wise

**Affiliations:** Department of Neurosciences, Imaging and Clinical Sciences, University “G. D’Annunzio” of Chieti-Pescara, Italy, Chieti, Italy; Institute for Advanced Biomedical Technologies, University “G. D’Annunzio” of Chieti-Pescara, Italy, Chieti, Italy; Research & Clinical Translation, Magnetic Resonance, Siemens Healthineers AG, Erlangen, Germany

**Keywords:** cerebrospinal fluid, hyperoxia, T_1_ mapping, magnetic resonance imaging, vascular permeability

## Abstract

**Background:** This study optimised a non-invasive magnetic resonance imaging technique to investigate the transfer of oxygen into cerebrospinal fluid (CSF) across the whole brain of healthy subjects.

**Methods:** A shortening of the T_1_ (longitudinal relaxation time) of CSF was induced by 100% hyperoxia and measured using a 3D SPACE sequence at 3T in 29 subjects, thereby demonstrating the diffusion of oxygen from blood to CSF. T_1_ mapping was performed at high resolution (0.9-mm isotropic, taking approximately 16 minutes) to capture anatomical details of the CSF spaces and was also repeated more rapidly at a lower resolution (3-mm isotropic, taking approximately 3.5 minutes) to capture the temporal dynamics of T_1_ changes.

**Results:** Significant region-dependent reductions in T_1_ were observed, indicating increased oxygen concentration in CSF. These occurred most prominently and rapidly in the cortical subarachnoid space and basilar cisterns, stabilising between 7 and 10 minutes after initiating hyperoxia, suggesting that oxygen diffusion primarily occurs via pial arteries and arteries at the base of the skull, both of which are in proximity to CSF-filled spaces. Over a timescale of 16 minutes, smaller T_1_ changes, only observed on the high-resolution T_1_ maps, occurred in the posterior lateral ventricles, where the choroid plexus is found, and the cisterna magna, possibly because of mixing effects with the adjacent basilar cisterns.

**Conclusions:** This study provides insights into the structure of CSF and dynamics of blood-CSF oxygen exchange, including their regional dependence. Moreover, the methodology presented here could, in the future, offer a valuable tool for characterising the passive diffusion of oxygen across cerebral blood vessel walls (i.e., their oxygen permeability), thereby providing a potential marker of cerebrovascular integrity.

## Background

The blood-brain barrier (BBB) and blood-CSF barrier (BCSFB) regulate the exchange of substances between blood, cerebrospinal fluid (CSF) and the adjacent brain parenchyma, playing a critical role in maintaining brain homeostasis and removing cerebral waste products. Changes in BBB and BCSFB permeability may reveal pathological alterations to these barriers, and characterising these exchange dynamics has become a growing focus of the magnetic resonance imaging (MRI) community [1].

In this context, practical methods are needed to investigate exchange dynamics between tissue compartments. Some studies on human patient populations have administered gadolinium-based contrast agents (GBCAs) either intrathecally [2–4] or intravenously [5–7] to characterise the time course of compartmental contrast enhancement. Intrathecal GBCA injection reveals the natural exit routes in the brain clearance system, whereas intravenous injection provides information on both entry and exit pathways. Intrathecally-injected GBCA caused enhancement observable on T_1_-weighted images within 20 minutes at the base of the skull (foramen magnum), within hours in the cortical sulci and cisternal spaces, and after 24 to 48 hours in brain tissue [2–4]. Intravenously-injected GBCA caused enhancement on T_2_-weighted fluid-attenuated inversion recovery (FLAIR) images with a different time course, as leakage into various CSF-filled regions started hours after injection [5–7]. However, these studies were conducted in clinical cohorts and their replication in healthy individuals is limited, due to the invasiveness of intrathecal GBCA injections [8] and growing concerns regarding gadolinium retention in brain tissue following intravenous administration [9].

Non-invasive techniques for CSF imaging have also been developed based on the transfer of water or molecular oxygen (O_2_) between tissue compartments. CSF production is thought to primarily occur in the cerebral ventricles at the BCSFB through the exchange of water and ions from the blood in the choroid plexuses [1]. To investigate the transfer of water from blood to CSF, some studies have used arterial spin labeling (ASL) with a long labeling duration and post-labeling delay, showing that labelled water reaches CSF from blood within seconds [10–12]. According to the two traditional models for brain fluid dynamics, namely the glymphatic system and the intramural peri-arterial drainage model (IPAD), water influx from the CSF into brain tissue occurs through specialised perivascular spaces and is driven by pulsatility of the arterial wall [1]. To investigate the transfer of water between CSF and brain parenchyma, one study has proposed an inversion-recovery approach with a long echo time (TE) [13] to measure the longitudinal relaxation time (T_1_) of CSF, reporting that shorter T_1_ in some regions could indicate faster CSF-brain tissue exchange.

Similarly to water, which passes through the BBB and BCSFB facilitated by aquaporin water channels and co-transporters, O_2_ can pass through the BBB and the BCSFB due to its lipid solubility and the high vascular permeability of vessels to O_2_ (particularly in the choroid plexus and pial vessels) in proximity to CSF [14]. This transport is ultimately driven by the large concentration gradient of O_2_ (a difference in oxygen partial pressure, ΔPO_2_, with PO_2_ indicating the amount of O_2_ physically dissolved in CSF) from the O_2_-rich arterial blood to CSF and tissue, and will depend on the vascular permeability to O_2_ [14]. Two studies have used custom inversion-recovery sequences to map the T_1_ of CSF before and during increased O_2_ inhalation (termed hyperoxia), as supplemental paramagnetic O_2_ shortens T_1_ of CSF relative to room-air breathing (termed normoxia), and thereby calculate changes in PO_2_ of CSF [15,16]. These studies have shown regional shortening of CSF T_1_ with hyperoxia and reported hyperoxia-induced effects in the CSF within minutes of O_2_ administration, demonstrating the transfer of O_2_ from the blood to the CSF compartment [15,16]. However, these [15,16] and other studies [17–22] performing CSF T_1_ mapping during hyperoxia [15–17] or normoxia [18–22] have been limited in spatial coverage, relying on stacks of thick imaging slices [15–20,22], or in image resolution, relying on 1-mm^2^ in-plane resolution or lower [15,17,19,22], or both. These limitations have precluded whole-brain coverage and, as only one study [16] supressed the signal from brain parenchyma, have generally rendered measurements susceptible to partial volume contamination from tissues adjacent to CSF.

Whole-brain coverage and sub-millimetric isotropic resolution are desirable in this context, as exchange dynamics between blood, CSF and brain tissue are likely to vary spatially across the brain and the ability to resolve smaller CSF-filled structures, such as enlarged perivascular spaces, may contribute to the understanding of brain fluid dynamics.

Here, we aimed to demonstrate non-invasively, with whole-brain coverage and sub-millimetric voxel resolution, the transfer of O_2_ between the arterial blood and CSF compartments in a healthy population. To this end, we measured T_1_ in the CSF using a tailored three-dimensional (3D) sampling perfection with application-optimised contrasts using different flip angle evolution (SPACE) pulse sequence [23], which uses non-selective, short refocussing pulse trains with varying flip angles less than 180 degrees. Compared to conventional fast/turbo spin echo, this combination reduces echo spacing while lengthening the usable duration of the spin-echo train, enabling high-resolution 3D spin-echo imaging with a lower specific absorption rate. A long effective TE (TE_eff_) provides a good suppression of signal from brain parenchyma while preserving signal from CSF given its considerably longer T_2_ [24]. To obtain information on the rate of O_2_ exchange between blood and CSF, we assessed changes in the CSF T_1_ during the transition from normoxia to hyperoxia (100% O_2_ administration). We obtained spatial T_1_ maps using a high-resolution version of the SPACE sequence to characterise the distribution of changes in PO_2_ across the full CSF compartment. In addition, we used a lower-resolution optimisation of the SPACE sequence for repeated T_1_ mapping to assess the temporal dynamics of O_2_ exchange during the transition from normoxia to hyperoxia.

## Methods

### Participants

MRI data were acquired in 29 healthy subjects (12/17 males/females, average age ± standard deviation: 35.7 ± 9.6 years, age range: 24-54 years). All participants reported no diagnosis of neurological, cardiovascular or respiratory conditions. This study was approved by the Institutional Review Board of the Department of Neurosciences, Imaging and Clinical Sciences, University of Chieti-Pescara (protocol number: 07/2023). All participants provided written informed consent and were instructed to abstain from consuming caffeinated or alcoholic beverages from the night before the experiment. This research was conducted in accordance with the World Medical Association Declaration of Helsinki.

### Experimental protocol

Before undergoing MRI, subjects were fitted with a breathing circuit for O_2_ delivery as described in Tancredi et al [25]. The tight-fitting mask was secured to the subject’s head using a four-point silicone harness. When the O_2_ supply is switched on, the presence of a reservoir ensures that the inspired O_2_ concentration remains consistently close to 100% even when the inspired gas flow during peak inspiration temporarily exceeds the O_2_ source supply rate; when the O_2_ supply is inactive, the circuit provides room air to the participant. The air composition inside the mask was monitored through a line connected to a Luer lock sampling port, which directed the samples outside the scanner room to a respiratory gas analyser (ML206, ADInstruments, Dunedin, New Zealand) connected to a PowerLab 16/35 data acquisition and analysis system (PL3516, ADInstruments). Samples were visualised and recorded using LabChart software (v8.1 Pro, ADInstruments). The subject’s heart rate and O_2_ saturation levels were continuously monitored using a wireless pulse oximeter (Expression MR200, Philips Healthcare, Best, The Netherlands).

All subjects underwent MRI on a 3 T system using a 32-channel receive-only head coil (MAGNETOM Prisma, Siemens Healthineers, Forchheim, Germany). At the beginning of the imaging session (Figure 1), structural images were acquired for anatomical reference using a magnetisation prepared 2 rapid acquisition gradient echoes (MP2RAGE) sequence [26] with the following parameters: sagittal slice orientation, phase encoding direction=anterior-posterior, resolution=1-mm isotropic, field of view (foot-head x anterior-posterior x right-left)=256x256x176 mm^3^, repetition time (TR)=5000 ms, TE=3.58 ms, inversion times (TIs)=700/2500 ms, flip angles=4/5 degrees, generalised autocalibrating partial parallel acquisition (GRAPPA) acceleration factor=3 [27], bandwidth=180 Hz/pixel and acquisition duration=4:59 min:s.

**Figure 1.**
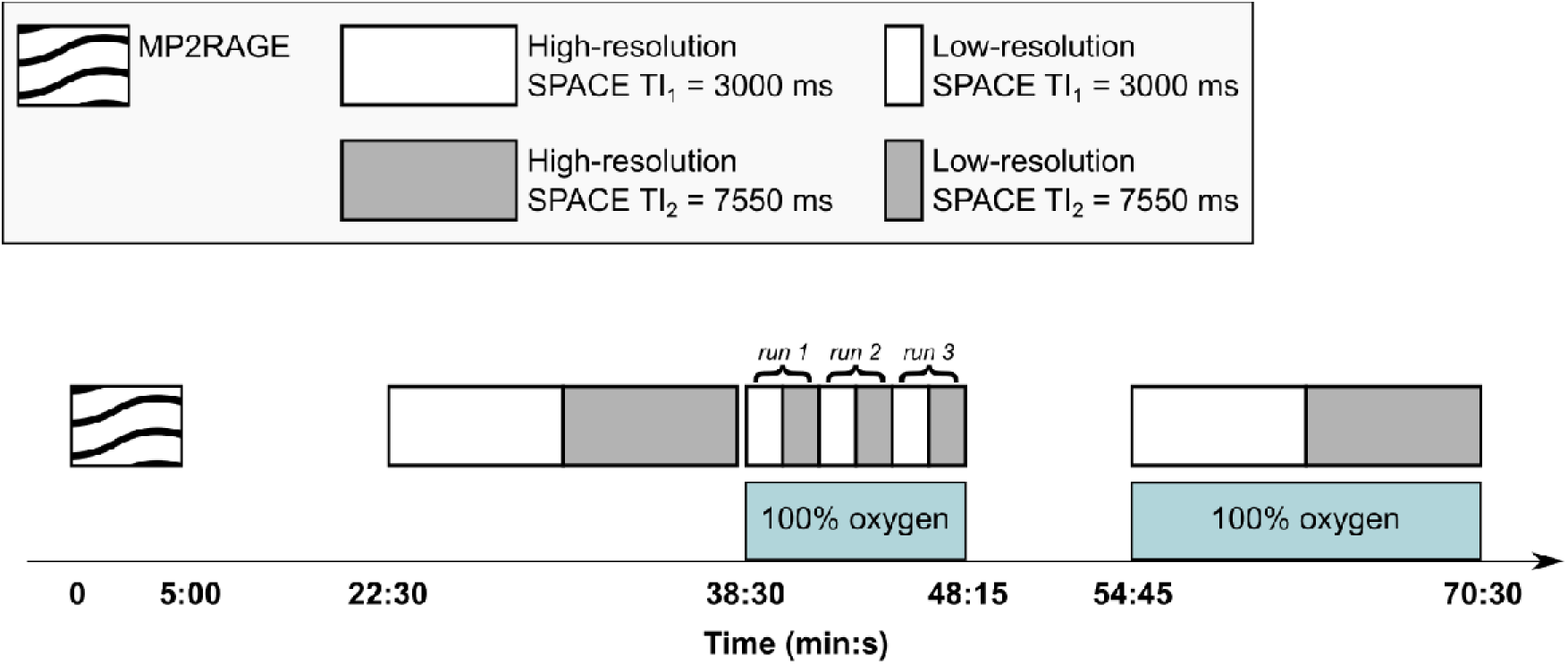
Experimental protocol. The diagram shows the time allocated for the sequences in this study. The region shaded in light blue corresponds to 100% O_2_ administration. The open time slots during the experiment were allocated for imaging unrelated to the present study. The labels ‘run 1’, ‘run 2’ and ‘run 3’ denote the three pairs (TI_1_, TI_2_) of low-resolution SPACE acquisitions performed during the first phase of O_2_ administration.

Two versions of the 3D SPACE sequence were implemented, one with an isotropic resolution of 0.9 mm and the other with an isotropic resolution of 3 mm. A schematic representation of the two SPACE sequence optimisations is shown in Supplementary Figure S1. Both employed a non-selective T_2_-preparation module (T_2,prep_=200 ms) followed by a long TE_eff_≍750 ms to exploit the markedly longer T_2_ of CSF relative to brain tissue. The two suppression mechanisms acted at distinct points in the sequence and are best described separately. The T_2_-preparation module attenuated the longitudinal magnetisation before the inversion pulse. With the chosen T_2_-preparation time, assuming an exponential T_2_ signal decay:

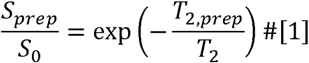

signals from the white and grey matter (T_2_≍80-110 ms [28]) retained 8-16% of their pre-preparation value, while signal from CSF (T_2_≍1.5-2.5 s [29]) retained 88-92% of its pre-preparation value.

During the subsequent inversion time, T_1_ recovery acted on the inverted magnetisation according to:

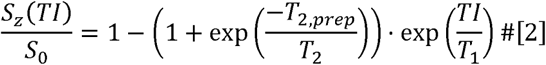

At TI_2_=7550 ms, which greatly exceeded the T_1_ of white matter (T_1_≍830 ms [28]) and grey matter (T_1_≍1330 ms [28]), parenchymal longitudinal magnetisation recovered to approximately 99-100% of S_0_, effectively erasing the effect of the T_2_-preparation attenuation step. At TI_1_=3000 ms, recovery was less complete, as white and grey matter respectively reached approximately 97% and 88% of S_0_. CSF (T_1_≍3.8-4.5 s [18–21]) recovered only partially during both TIs, retaining the influence of the T_2_-preparation module.

The subsequent long TE_eff_ provided T_2_ weighting at signal readout, acting on this recovered longitudinal magnetisation:

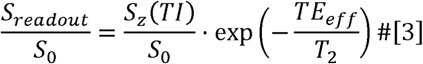

For brain parenchyma, where T_1_ recovery during TI restored S_z_ to approximately S_0_, Equation 3 simplified to:

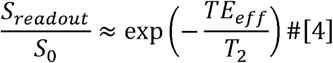

reducing the parenchymal signal by approximately three orders of magnitude and leaving a residual of 0.008-0.11% of S_0_ at both TIs. For CSF, combining Equations 2 and 3 across the expected T_1_ and T_2_ ranges [18–21,29] yielded a signal at readout of approximately 39-55% of S₀ at TI_2_, where CSF signal neared its recovery plateau. At TI_1_, CSF signal at readout was intentionally reduced to approximately 1-11% of S₀, as this acquisition was positioned near the CSF null point to maximise the dynamic range of the TI_2_/TI_1_ signal ratio. The residual signal from brain parenchyma at the time of k-space centre acquisition was therefore negligible relative to the CSF signal.

To measure T_1_, the high-resolution implementation of the 3D SPACE sequence consisted of two separate acquisitions, with all parameters held constant except for the TIs, which were set to 3 s (TI_1_) and 7.55 s (TI_2_) in the first and second acquisition, respectively. The other parameters were: sagittal slice orientation, phase encoding direction=anterior-posterior, field of view (foot-head x anterior-posterior x right-left)=230x230x158 mm^3^, 0.9-mm isotropic resolution, TR=10010 ms, magnitude preparation=non-selective T_2_ preparation inversion recovery, T_2,prep_=200 ms, TE_eff_=751 ms, turbo factor=599, refocussing flip angle=120 degrees, flip angle mode=constant, wrap-up magnitude=RESTORE, GRAPPA acceleration factor=2, bandwidth=888 Hz/pixel and acquisition duration=7:52 min:s. A constant refocussing flip angle below 180 degrees was chosen because, unlike variable flip angle trains, it produces a signal evolution governed by a predictable interplay of T_1_ and T_2_ decay (including stimulated echo contributions) that can be explicitly characterised and fully accounted for in Bloch equation simulations [24]. Thus, the time required to obtain a high-resolution T_1_ map was approximately 16 minutes.

The low-resolution implementation of the 3D SPACE sequence also consisted of two separate acquisitions and had similar parameters as the high-resolution version, except for the following: resolution=3-mm isotropic, field of view (foot-head x anterior-posterior x right-left)=384x306x216 mm^3^, TE_eff_=752 ms, flip angle=115 degrees, bandwidth=494 Hz/pixel, and acquisition duration=1:42 min:s. Thus, the time required to obtain a low-resolution T_1_ map was approximately 3.5 minutes.

The TI values of the SPACE sequence were chosen based on the expected T_1_ of CSF at 3T, which has most frequently been reported in the range of 3.8-4.5 s [18–21] (with just one study reporting a much higher average T_1_ of 6.9 s in three female subjects [20]). TI_1_=3 s was set near the CSF null point (Tl_nuU_=T_1_.In 2≈ 2.6-3.1s), where fractional signal changes are most pronounced and T_1_ sensitivity is high, while TI_2_=7.55 s was placed near the recovery plateau to maximise the dynamic range of the signal ratio. The TR was chosen to allow sufficient longitudinal recovery of CSF between repetitions, aided by the RESTORE module, which returns residual transverse magnetisation to the longitudinal axis at the end of each echo train (Supplementary Figure S1).

The high-resolution sequence was employed to calculate T_1_ maps at baseline during normoxia and at the end of the experiment during hyperoxia (Figure 1, second phase of O_2_ administration). The low-resolution sequence, with a shorter acquisition time, was used to calculate three T_1_ maps during the first phase of O_2_ administration, providing information on the temporal dynamics of CSF T_1_ changes caused by hyperoxia. The repeats of each pair of low-resolution acquisitions (TI_1_, TI_2_) will be referred to as runs 1, 2 and 3 (Figure 1). Notably, because it was acquired after the start of O_2_ administration, run 1 could not be considered a true baseline for T_1_ values at a low resolution. Thus, an exploratory analysis of baseline T_1_ at a low resolution was performed in a subset of four subjects by adding an additional low-resolution SPACE acquisition at normoxia (before the first phase of O_2_ administration).

O_2_ was administered in two phases of the experiment (Figure 1) at a flow rate of 15 L/min and humidification was provided for subject comfort. Initially, 100% O_2_ was administered for 10 minutes. This duration has been shown to yield the maximum observable change in CSF T_1_ during 100% hyperoxia [15]. Thus, acquiring three runs enabled the calculation of T_1_ maps to investigate dynamic changes in T_1_. Then, O_2_ administration was paused for approximately 6 minutes and 30 seconds. During this period, a sequence unrelated to the present study was conducted. This pause was inserted to adhere to local safety guidelines regarding 100% O_2_ administration. Afterwards, 100% O_2_ administration was resumed until the end of the experiment, as shown in Figure 1. The effects of hyperoxia on CSF were expected to return to the pre-pause level within the duration of the second acquisition of the high-resolution SPACE TI_1_ sequence (Figure 1) [15].

### CSF T_1_ mapping

Image analysis was performed using MATLAB (R2022b, The MathWorks, Natick, MA), the FMRIB software library (FSL) v6.0 [30], and the Advanced Normalisation Tools (ANTs) software library [31]. All rigid image coregistrations within subject space were performed using FSL FLIRT [32,33], chosen for its speed and accuracy comparable to ANTs for rigid registration. Non-linear coregistrations of structural MP2RAGE images to standard space were performed using the ANTs SyN function, which was chosen for its high accuracy [34]. Flowcharts of the image analysis steps for CSF T_1_ mapping are shown in Supplementary Figure S2.

The first step in the pipeline for T_1_ mapping was to ensure image alignment within and between conditions. For each pair of high-resolution SPACE acquisitions, the TI_2_ image was rigidly coregistered to the TI_1_ image. Moreover, to align normoxia and hyperoxia acquisitions, the high-resolution TI_1_ image at hyperoxia was rigidly coregistered to the corresponding normoxia TI_1_ image. Similarly, within-pair rigid coregistration was applied independently to each run of the low-resolution SPACE acquisition. Where multiple sequential transformations were required, transformation matrices were combined prior to resampling to minimise the number of interpolation steps and reduce associated interpolation errors.

A high-resolution mask of CSF was then calculated based on the difference between the aligned high-resolution TI_2_ and TI_1_ hyperoxia images, thereby exploiting the higher signal-to-noise ratio resulting from breathing 100% O_2_ compared to breathing room air. To remove any residual tissue signal, the difference between TI_2_ and TI_1_ was empirically thresholded at 50% of its robust range, that is, the intensity distribution with outlier values removed, and then binarised. MATLAB’s *regionprops* function was applied to select the largest connected component from the binary mask. The resulting CSF mask was then rigidly transformed into high-resolution normoxia space by applying the previously calculated hyperoxia-to-normoxia transformation. For each run of the low-resolution SPACE sequence, the CSF mask was propagated from the high-resolution hyperoxia acquisition by rigidly aligning the high-resolution hyperoxia TI_1_ image with the low-resolution TI_1_ image of each run independently, and applying the resulting transformation to the high-resolution CSF mask.

Finally, for each voxel within the CSF mask and each pair of TI acquisitions, the measured TI_2_/TI_1_ signal ratio was matched to a Bloch-simulation lookup table (LUT) to estimate apparent T_1_. For each SPACE optimisation, the LUT was generated over expected CSF T_1_ and T_2_ ranges at 3 T [18–21,29] (T_1_ range: 3000-6500 ms, step size=50 ms; T_2_ range: 1500-2500 ms; step size=10 ms) by simulating the full inversion-prepared, variable-flip-angle SPACE echo train (including T_2_-preparation, inversion, echo spacing, flip-angle series and RESTORE) for each (T_1_, T_2_) pair and sampling the signal at the central k-space line. Because the resulting TI_2_/TI_1_ ratio is a smooth, monotonic function of T_1_ and effectively T_2_-independent over the simulated CSF range, T_1_ was estimated by a well-conditioned one-dimensional matching, and the LUT-based T_1_ estimation was considered effectively T_2_-insensitive.

Notably, where additional low-resolution SPACE acquisitions at normoxia were available (four subjects), all analysis steps (coregistration, mask propagation, and T_1_ mapping) were applied identically to those runs.

### T_1_ map alignment and difference image calculation

For visual inspection of T_1_ alterations induced by hyperoxia, the high-resolution T_1_ maps calculated at hyperoxia and normoxia were rigidly aligned based on the transformation between the respective TI_1_ images calculated earlier. Similarly, for visual inspection of the dynamic effect of O_2_ inhalation on T_1_, all low-resolution T_1_ maps were aligned in the space of run 1 by calculating the rigid transformations between the TI_1_ images from runs 2 and 3 and the TI_1_ image from run 1, and applying the resulting transformations to the corresponding T_1_ maps. Then, difference images were computed between the high-resolution T_1_ maps obtained at normoxia and hyperoxia, and between each pair of low-resolution T_1_ maps.

### Regional analysis of T_1_ values

For regional analysis of T_1_ values in the CSF, the following regions of interest (ROIs) were manually delineated using ITK-SNAP within the contours of the high-resolution CSF mask: the anterior lateral ventricles (anterior to the foramen of Monro), the posterior lateral ventricles (posterior to the foramen of Monro, including the body, atrium, occipital and temporal horns of the lateral ventricles), the third ventricle (inferior to the foramen of Monro and superior to the cerebral aqueduct), the cortical subarachnoid space, the infratentorial basilar cisterns (encompassing the interpeduncular, the prepontine and premedullary cisterns), the fourth ventricle and the cisterna magna. These ROIs, encompassing CSF-filled regions in the whole brain, were selected to enable comparison with previous studies investigating the T_1_ values of CSF during hyperoxia [15,17] and a study suggesting that a faster CSF-tissue exchange rate characterises regions with lower T_1_ values [13]. For each subject and all high-resolution and low-resolution T_1_ maps, the median T_1_ value was calculated in each ROI.

Statistical analyses of the regional median values of T_1_ were performed using JASP (v0.97.1) [35]. All statistical tests were two-tailed, and *p*<0.05 was considered significant.

A two-way repeated-measures analysis of variance (ANOVA) was performed to test for differences in regional T_1_ values between normoxia and hyperoxia conditions based on the high-resolution SPACE data. Two repeated-measures factors were set: condition (normoxia or hyperoxia) and ROI. To account for potential differences in CSF clearance influenced by the age and sex of the participants [36,37], age was included as a covariate and sex as a between-subjects factor.

A second two-way repeated-measures ANOVA was performed to assess whether hyperoxia-induced changes in CSF T_1_ stabilise over the couse of the hyperoxic challenge, and whether any temporal dynamics differed across ROIs, based on the low-resolution SPACE data. Two repeated-measures factors were set: time of acquisition (Time, runs 1 to 3) and ROI. Age was included as a covariate and sex as a between-subjects factor.

For both ANOVAs, assumptions were checked as follows: Mauchly’s test evaluated sphericity for factors with more than two levels, and, when violated, degrees of freedom were corrected using the Greenhouse-Geisser method; Q-Q plots visually assessed normality of residuals; and Levene’s test evaluated homogeneity of variances. All post hoc analyses were Holm-corrected for multiple comparisons.

### Analysis in standard space

Higher-level analysis of T_1_ differences between normoxia and hyperoxia was performed to identify O_2_-induced T_1_ shortening at the group level. To this aim, for each subject, both the high-resolution T_1_ maps calculated at normoxia and hyperoxia were registered to the 1-mm isotropic Montreal Neurological Institute (MNI) template [38,39] as follows.

First, the TI_1_ image of each SPACE acquisition was rigidly aligned to the structural T_1_-weighted MP2RAGE uniform (UNI) image. Compared to conventional T_1_-weighted sequences at 3 T, the MP2RAGE UNI image is designed to provide uniform intensity across the scanned volume by correcting for non-T_1_ contrast effects, such as receiver field (B_1_) inhomogeneity, proton density contrast and transverse decay (T_2_*) [26]. Moreover, it presents no signal in the CSF, thereby providing a contrast complementary to that observed in the SPACE acquisitions. To enable more accurate registration, strong background noise in air-filled regions was removed from the MP2RAGE UNI image using the intensity-normalised MP2RAGE image from the second inversion time (INV2) as a pseudo-mask via voxelwise multiplication (MPRAGEise v2.0, github.com/srikash/MPRAGEise). The denoised MP2RAGE UNI image was then bias field corrected (N4biasfieldcorrection, ANTs) [40]. Brain segmentation was achieved using the Computational Anatomy Toolbox (CAT) [41] implemented in the Statistical Parametric Mapping suite (SPM12; fil.ion.ucl.ac.uk/spm) for MATLAB, and a brain mask was obtained by combining the white and grey matter segmentations. The extracted brain region was then coregistered to the 1-mm isotropic MNI template using ANTs. The same image transformation was applied to align the individual high-resolution T_1_ maps to MNI space.

Using a custom implementation of nearest-neighbour interpolation in MATLAB, each T_1_ map in MNI space was downsampled to an isotropic 3-mm resolution. This downsampled voxel size was chosen to reduce the impact of slight misalignment between individual T_1_ maps in MNI space and improve the statistical power of group-level analysis. Because the T_1_ maps of CSF were sparse in nature, with NaN values assigned to voxels with no CSF signal, the custom implementation of nearest-neighbour interpolation explicitly checked for NaN values at the nearest-neighbour location. If a NaN value was found, the algorithm searched a 3x3x3 neighbourhood for the closest non-NaN value to fill the gap in the downsampled image.

The average T_1_ maps at normoxia and hyperoxia were calculated across subjects for display purposes. To account for data sparsity, the average T_1_ value of a voxel was only calculated if the voxel had a value different from NaN in at least 90% of subjects. Otherwise, the average of T_1_ in that voxel was set to zero. To assess differences in T_1_ of CSF between normoxia and hyperoxia, non-parametric permutation tests were conducted using the *randomise* tool in FSL [42]. Two distinct one-tailed one-sample t-tests were performed to test for ΔT_1_ > 0 and ΔT_1_<0 (where Δ denotes a difference) by providing as input a series of difference images (one per subject) between the T_1_ map calculated at normoxia and the T_1_ map calculated at hyperoxia. To maintain the family-wise error rate at 0.05, a threshold of *p*<0.025 was used for each of these two tests (Bonferroni correction). The binary CSF mask for the tests was calculated to include the voxels appearing in both the average T_1_ maps at normoxia and hyperoxia. For each test, 5000 permutations were used, along with the bi-dimensional version of threshold-free cluster enhancement [43], which is designed to correct for multiple comparisons when using sparse images.

## Results

### Overall CSF appearance on SPACE acquisitions

Figure 2 shows images acquired using the high-resolution SPACE sequence (Figures 2a-b) and the low-resolution SPACE sequence (Figures 2c-d) at the two TIs. Lower windowing was applied to the images at the shorter TI, where CSF was nearly null (Figures 2a and 2c). Notably, residual non-CSF signal appeared only outside the brain, suggesting that the chosen T_2_ preparation and the effective TE minimised signal contributions from brain parenchyma (Figure 2).

**Figure 2.**
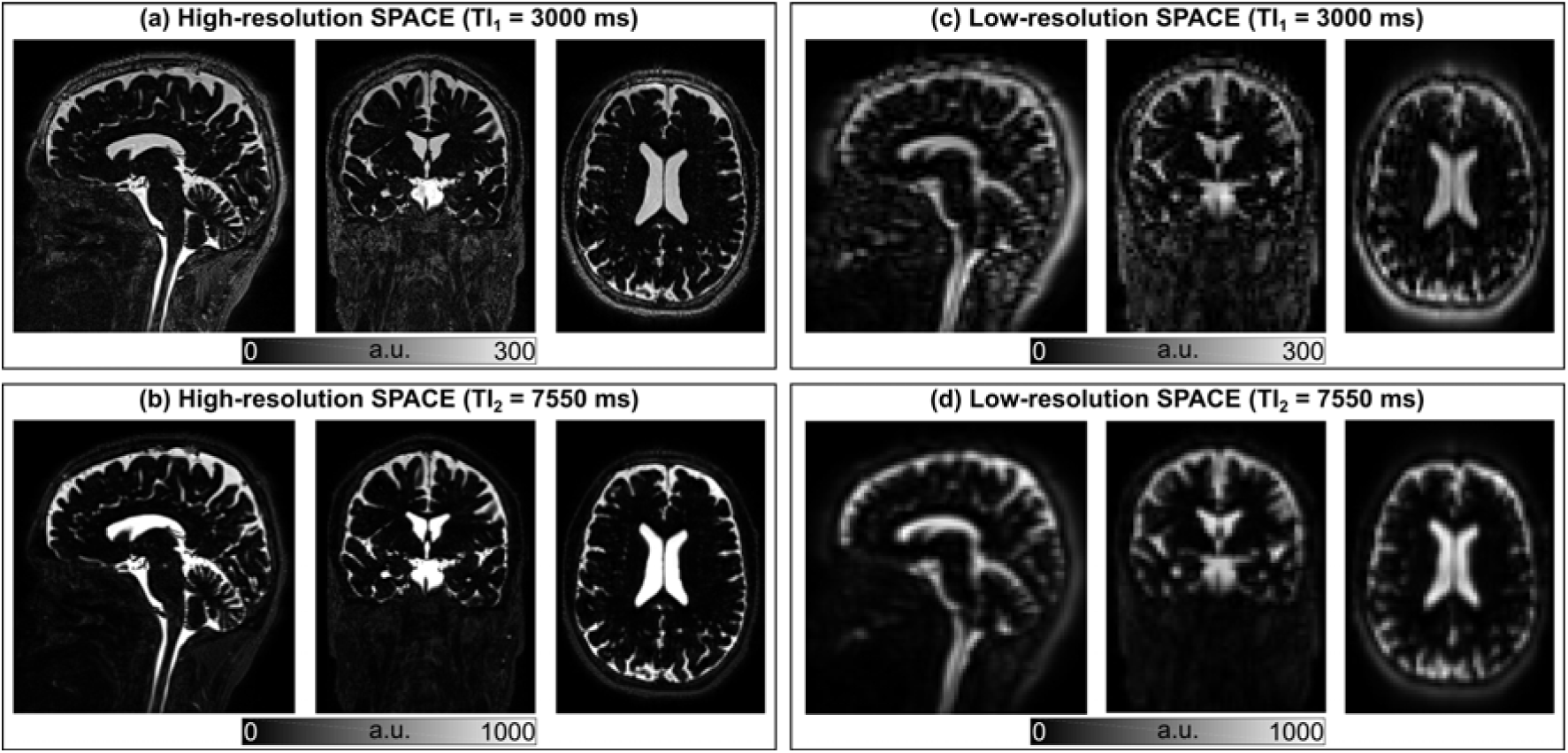
Representative SPACE acquisitions. For a representative subject, the same sagittal, coronal and transverse slices are shown for the high-resolution SPACE acquisition acquired at normoxia at TI_1_ (a) and TI_2_ (b). Similar slices are shown for the first run (at hyperoxia) of the low-resolution SPACE acquisition at TI_1_ (c) and TI_2_ (d). The slices depicted in (c) and (d) were cropped to match the field of view of those in (a) and (b). Note that different display windows were applied to the TI_1_ and TI_2_ images; the scale bars in each panel indicate the corresponding signal intensity range, reflecting the substantially lower CSF signal at TI_1_ relative to TI_2_.

### T_1_ mapping

Figure 3 shows representative high-resolution T_1_ maps acquired at normoxia (Figure 3a) and hyperoxia (Figure 3b). It also shows the difference between these two maps (Figure 3c) and the ROIs used for regional analysis of T_1_ (Figure 3d). Compared to breathing room air, O_2_ inhalation caused widespread T_1_ shortening in the cortical subarachnoid space (Figures 3a-c). Similarly, Figure 4 shows representative low-resolution T_1_ maps acquired at hyperoxia at each run (Figures 4a-c) and the T_1_ difference map between each pair of runs (Figures 4d-f).

**Figure 3.**
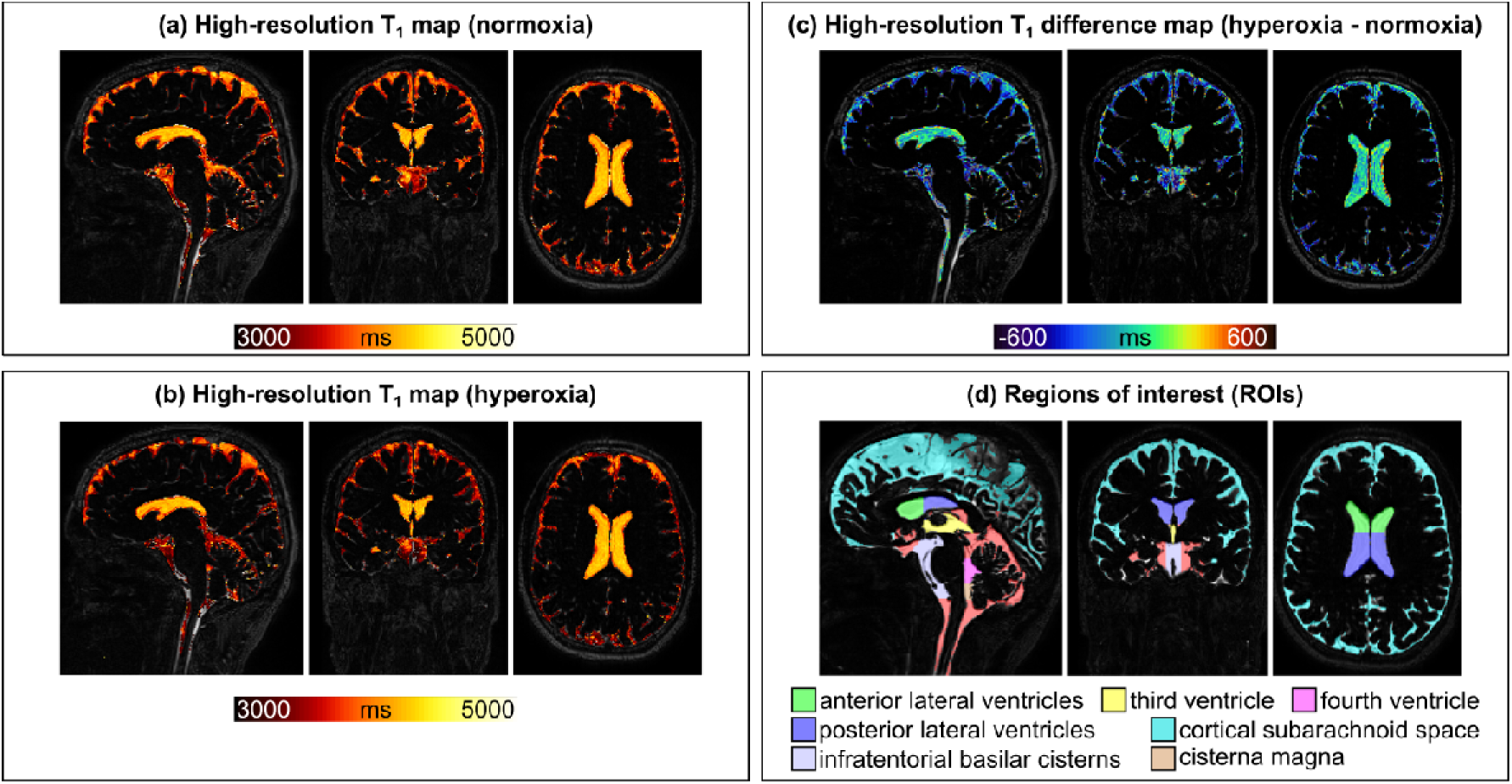
High-resolution T_1_ maps. For the same representative subject as in Figure 2, the same sagittal, coronal and transverse slices are shown for the T_1_ maps acquired during normoxia (a), hyperoxia (b) and the difference between the two (c). The maps in panels a-c are overlaid on the high-resolution SPACE image acquired at normoxia at TI_1_. The ROIs used for regional analysis of T_1_ values in the CSF are shown in (d).

**Figure 4.**
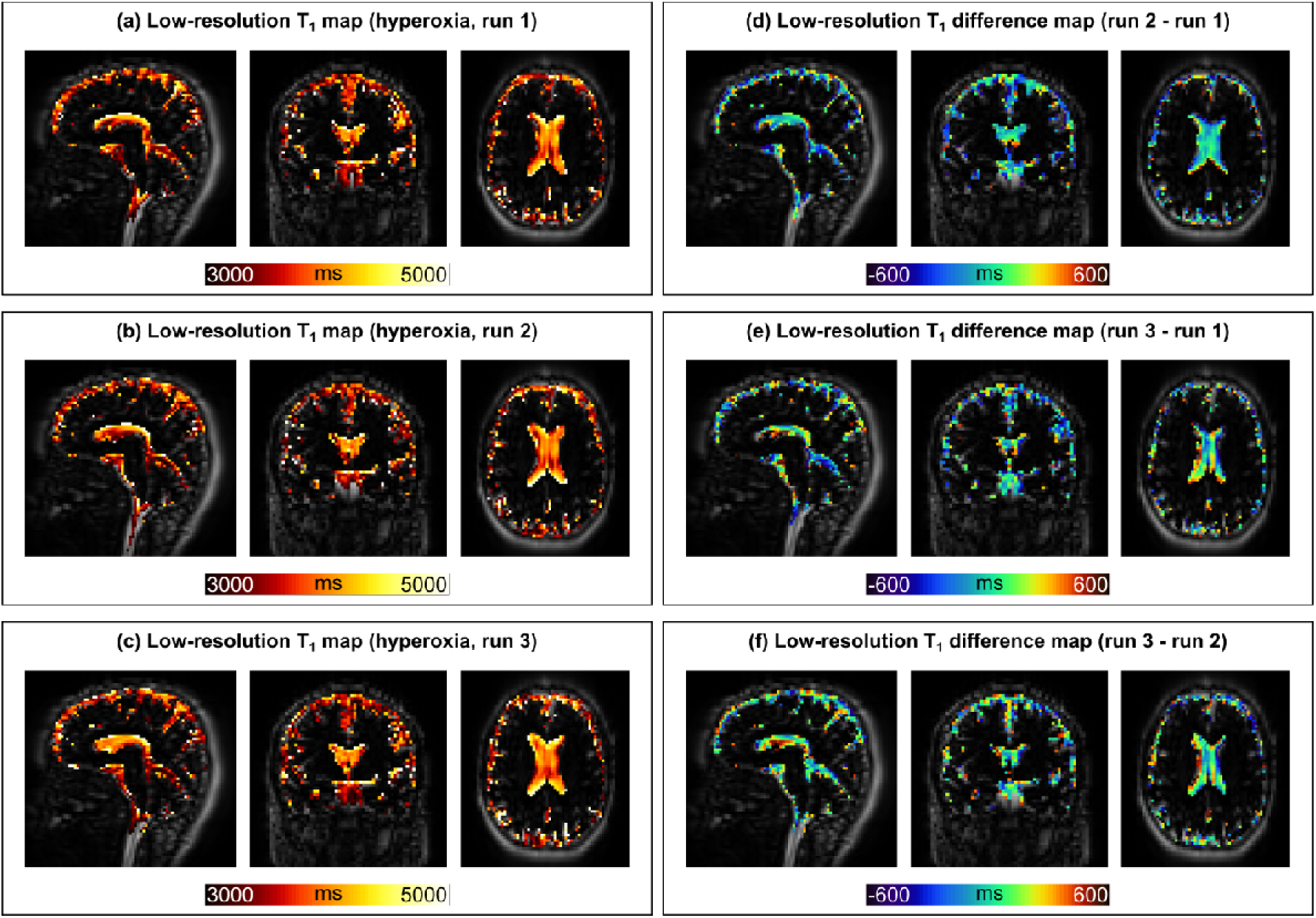
Low-resolution T_1_ maps. For the same representative subject as in Figure 2, the same sagittal, coronal and transverse slices are shown for the T_1_ maps acquired during hyperoxia at run 1 (a), run 2 (b), and run 3 (c). T_1_ difference maps between each pair of runs are shown in (d-f). All maps are overlaid on the low-resolution SPACE image acquired at run 1 at TI_1_.

### Regional analysis of T_1_

The regional values of T_1_ are shown using raincloud plots for both the high-resolution (Figure 5) and low-resolution SPACE acquisitions (Figure 6). Raincloud plots for the four subjects that also underwent a baseline low-resolution SPACE acquisition at normoxia are shown in Supplementary Figure S3.

**Figure 5.**
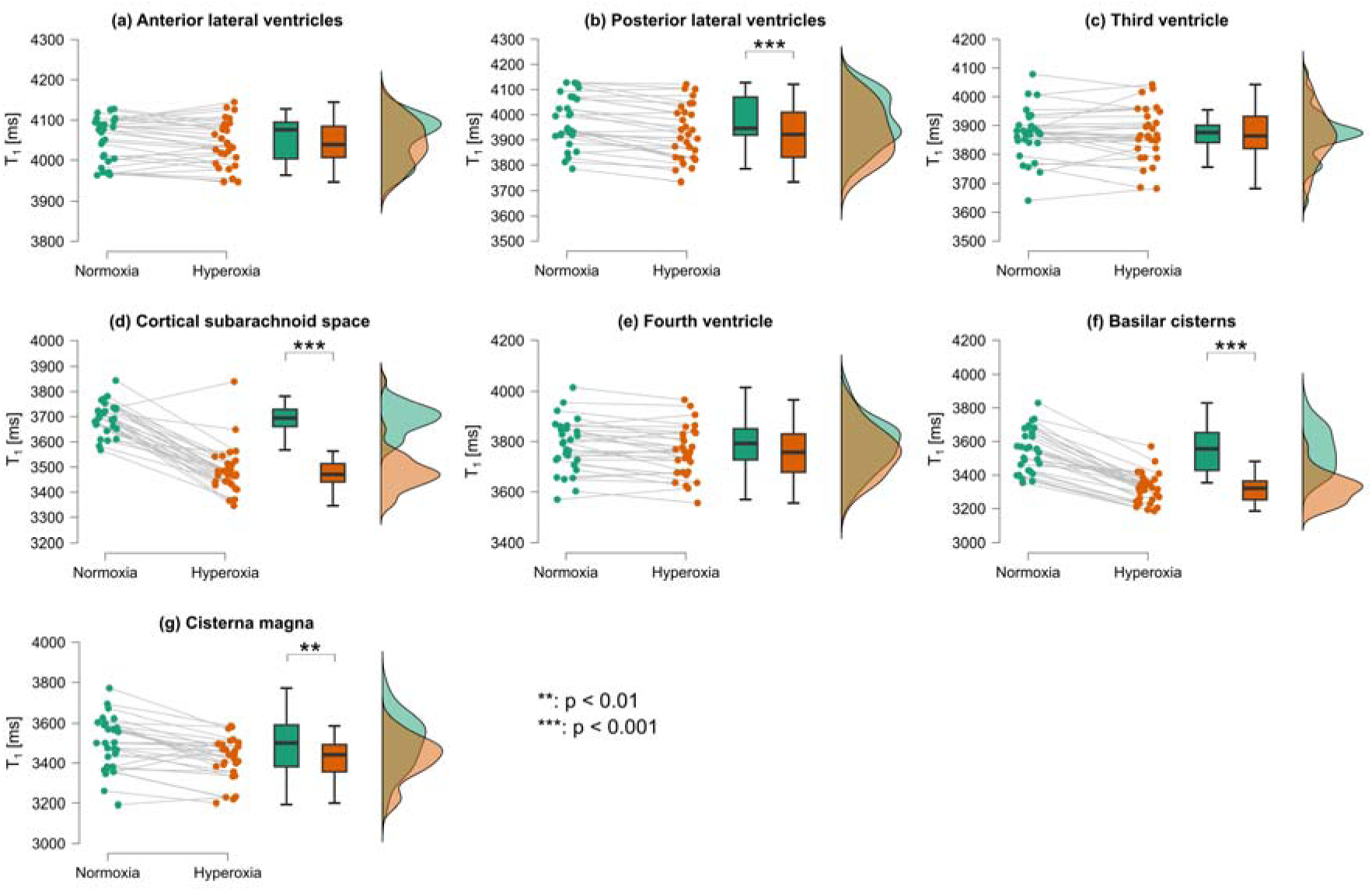
Raincloud plots of regional high-resolution T_1_ measurements. For each ROI, the individual subject median values of T_1_ measured during normoxia and hyperoxia are shown using a raincloud plot. This plot includes, for each experimental condition, a scatter plot of individual data points, a box plot, and a histogram. In the scatter plots for each region, pairs of data points connected by a line indicate measurements from the same subject under the two experimental conditions. Significant differences between the two experimental conditions are denoted as ** p<0.01, *** p<0.001.

**Figure 6.**
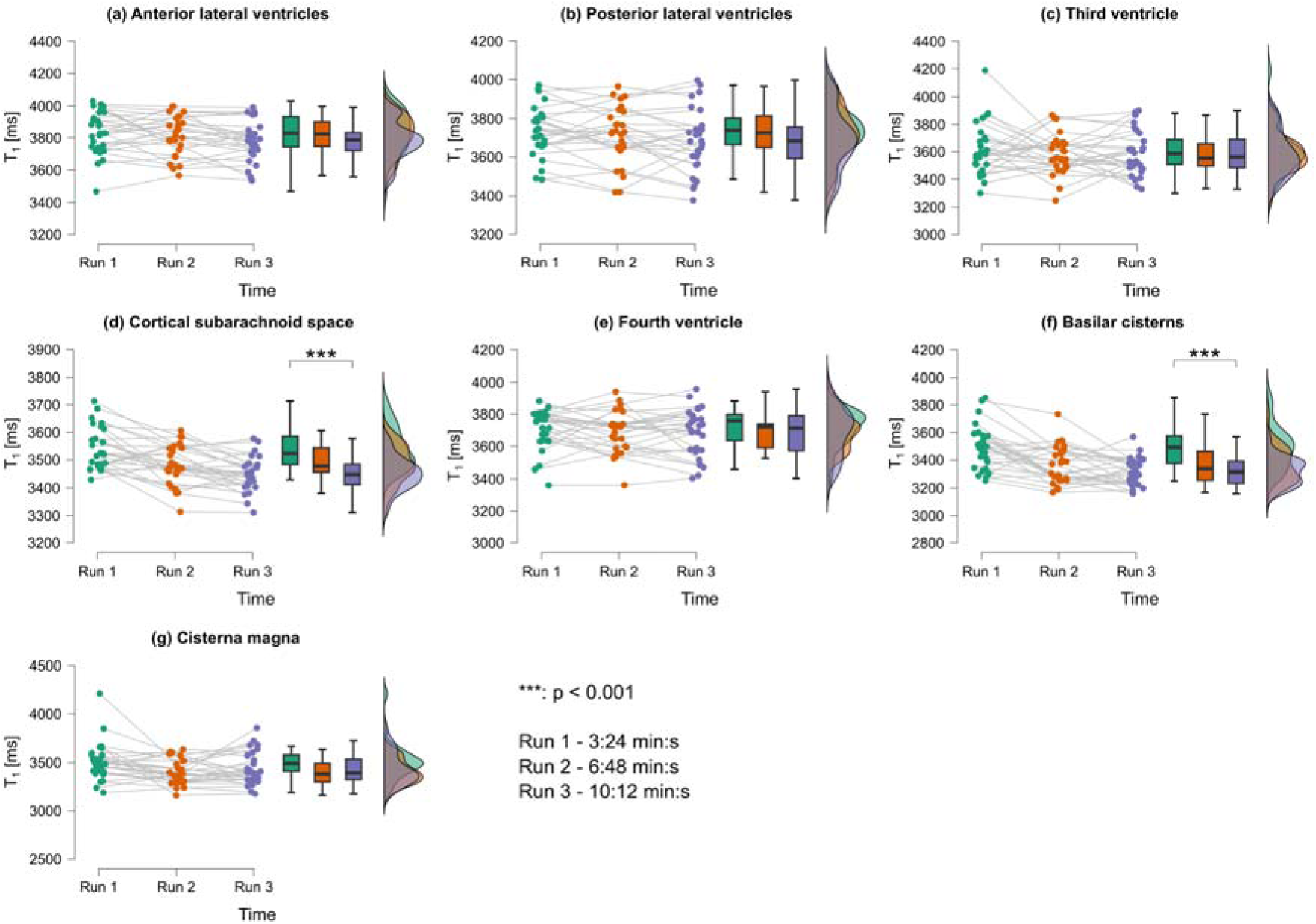
Raincloud plots of regional low-resolution T_1_ measurements. For each ROI, the individual median values of T_1_ measured at three time points (runs 1, 2 and 3) during hyperoxia are shown using a raincloud plot. This plot includes, for each run, a scatter plot of individual data points, a box plot, and a histogram. In the scatter plots for each region, triplets of data points connected by a line indicate measurements from the same subject taken at each time point. Significant differences between the three time points are denoted as *** p<0.001.

In both repeated measures ANOVA tests, Mauchly’s test of sphericity was significant, so the Greenhouse-Geisser correction was applied. The assumptions of normality and variance homogeneity were satisfied.

#### Regional high-resolution T_1_ measurements

T_1_ of CSF differed significantly across ROIs, *F*(3.81, 99.02)=19.71, *p*<0.001, ω^2^=0.298, and effect of hyperoxia on T_1_ was ROI-dependent, *F*(3.76, 97.83)=3.98, *p*=0.006, ω^2^=0.023.

Across all ROIs, hyperoxia was associated with a significant overall reduction in T_1_ compared to normoxia (mean difference=82.1 ms, standard error (SE)=6.2 ms, *t*(26)=13.22, *p*<0.001). Post hoc comparisons revealed that this reduction was significant in the posterior lateral ventricles (mean difference=41.9 ms, SE=7.5 ms, *p*<0.001; Figure 5b), cortical subarachnoid space (mean difference=201.9 ms, SE=17.1 ms, *p*<0.001; Figure 5d), basilar cisterns (mean difference=223.2 ms, SE=17.3 ms, *p*<0.001; Figure 5f), and cisterna magna (mean difference=77.1 ms, SE=17.4 ms, *p*=0.002; Figure 5g), but was not significant in the anterior lateral ventricles (mean difference=7.8 ms, SE=5.8 ms, *p*=0.567; Figure 5a) or third ventricle (mean difference=-1.3 ms, SE=8.3 ms, *p*=1.000; Figure 5c). The reduction in the fourth ventricle did not reach significance (mean difference=24.2 ms, SE=8.6 ms, *p*=0.055; Figure 5e).

Pairwise comparisons across ROIs (averaged across normoxia and hyperoxia) demonstrated significant differences between almost all region pairs (all *p*<0.001), with the single exception of the basilar cisterns versus the cisterna magna (mean difference=-23.3 ms, SE=20.0 ms, *p*=0.256). T_1_ values were longest in the lateral ventricular regions (anterior and posterior lateral ventricles) and shortest in the posterior fossa cisterns (basilar cisterns and cisterna magna).

#### Regional low-resolution T_1_ measurements

T_1_ of CSF differed significantly across ROIs (*F*(4.11, 102.79)=5.42, *p*<0.001, ω²=0.093), consistent with the regional pattern observed in the high-resolution data. Neither the main effect of Time (*F*(1.65, 41.25)=1.99, *p*=0.157, ω²=0.010) nor the Time × ROI interaction (*F*(5.57, 139.17)=0.76, *p*=0.596, ω²=0.000) reached significance, indicating that T_1_ did not change significantly across the three successive runs during hyperoxia. Neither sex nor age exerted a significant main effect (both *p*>0.05), nor did either interact significantly with Time or ROI (all *p*>0.05, all ω²≈0).

Post hoc comparisons of Time (averaged across ROIs) indicated that T_1_ values at run 1 were significantly higher than at both run 2 (mean difference=48.9 ms, SE=16.2 ms, *t*(25)=3.03, *p*=0.011) and run 3 (mean difference=66.4 ms, SE=10.5 ms, *t*(25)=6.33, *p*<0.001), while runs 2 and 3 did not differ significantly (mean difference=17.5 ms, SE=14.1 ms, *p*=0.227).

Pairwise comparisons within each ROI across time points revealed that this temporal pattern was specific to the cortical subarachnoid space and basilar cisterns. In the cortical subarachnoid space, T_1_ was significantly lower at run 3 than at run 1 (mean difference=88.8 ms, SE=13.2 ms, *p*<0.001; Figure 6d), while runs 2 and 3 did not differ significantly (*p*=0.130). In the basilar cisterns, T_1_ was significantly lower at run 3 than at run 1 (mean difference=162.5 ms, SE=31.4 ms, *p*=0.002; Figure 6f), while runs 2 and 3 again did not differ significantly (*p*=1.000). No significant differences across runs were observed in any other ROI (all *p*>0.05; Figures 6a-c, 6e, 6g). This suggests that the hyperoxia-induced T_1_ reduction in the cortical subarachnoid space and basilar cisterns continues to develop over the first ∼10 minutes of hyperoxia, approaching stability between approximately 7 and 10 min, whereas T_1_ values in the remaining regions did not change significantly across the duration of the acquisition.

Pairwise comparisons across ROIs (averaged across runs) demonstrated significant differences between almost all region pairs (all *p*<0.001), with the exceptions of posterior lateral ventricles versus fourth ventricle (*p*=0.304), cortical subarachnoid space versus cisterna magna (*p*=0.153), and basilar cisterns versus cisterna magna (*p*=0.153), broadly consistent with the high-resolution findings.

In the four subjects for which low-resolution T_1_ values were also calculated at normoxia, no statistical analysis was performed due to the limited sample size. However, visual inspection of the raincloud plots (Supplementary Figure S3) revealed that T_1_ values remained stable between normoxia and all runs at hyperoxia in both the anterior and posterior lateral ventricles and the third ventricle (Supplementary Figures S3a-c). In all other regions, qualitative T_1_ shortening between normoxia and run 1 at hyperoxia could be detected (Supplementary Figures S3d-e and S3g), with the exception of the basilar cisterns where T_1_ appeared to increase between normoxia and run 1 at hyperoxia (Supplementary Figure S3f).

### Higher-level analysis of T_1_

Figure 7 displays, in MNI space, the across-subject average of the high-resolution T_1_ maps of CSF acquired during normoxia (Figure 7a) and hyperoxia (Figure 7b), along with the T_1_ difference map between these two conditions (Figure 7c) and the (1-*p*) voxel-wise statistical map resulting from the permutation test assessing higher T_1_ values at normoxia than hyperoxia (Figure 7d). Results from the permutation test assessing lower T_1_ values at normoxia than hyperoxia are not shown, as no significant differences were found. The group-level analysis revealed significant T_1_ shortening widespread across the cortical subarachnoid space, the basilar cisterns, and, to a lesser degree, in posterior ventricular regions (Figure 7d). T_1_ values appeared unchanged in the anterior ventricles (Figure 7d).

**Figure 7.**
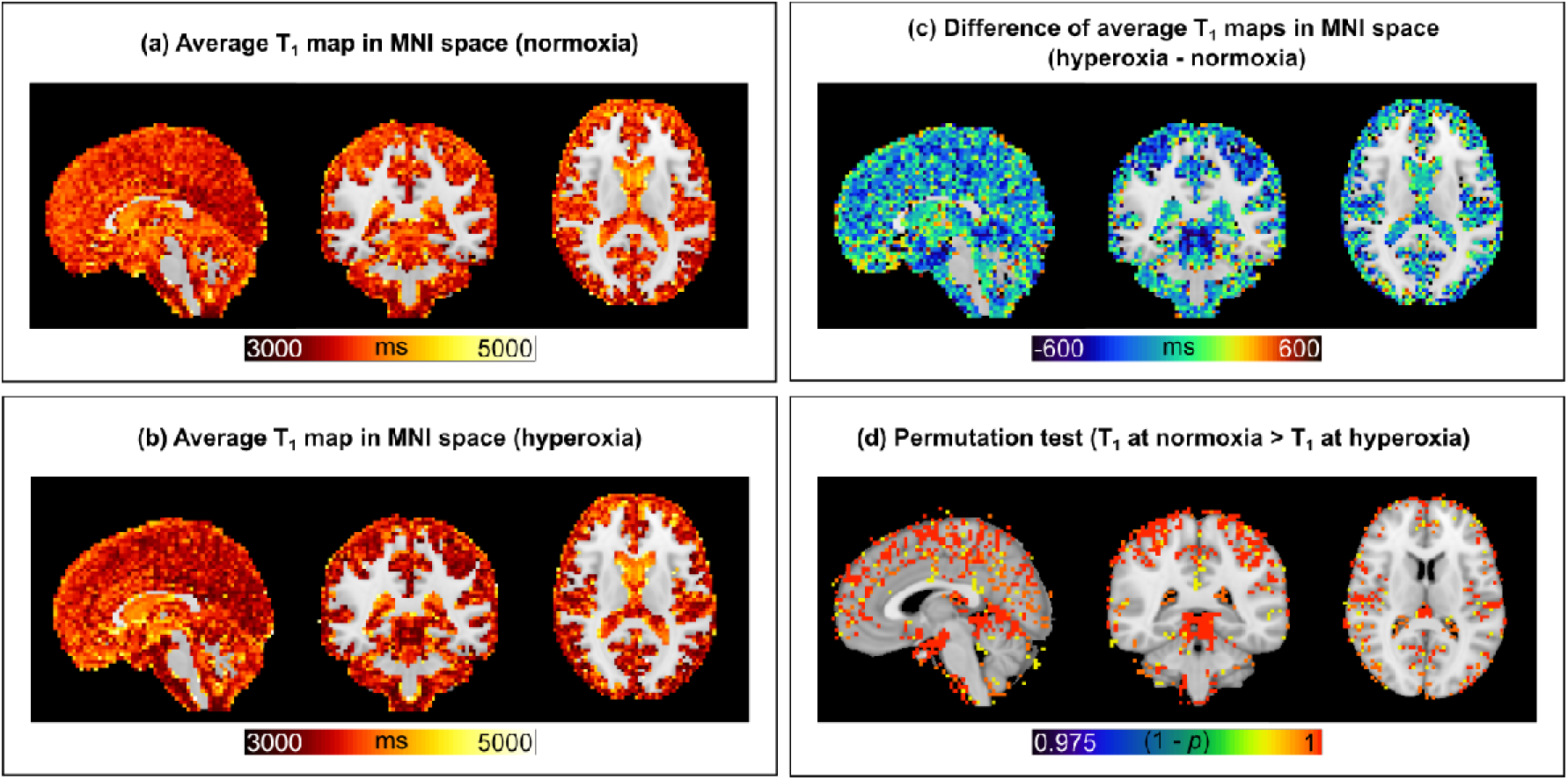
Average T_1_ maps in MNI space. The same sagittal, coronal and transverse slices are shown for the average T_1_ maps in MNI space acquired during normoxia (a), hyperoxia (b) and the difference between the two (c). The results of the non-parametric permutation test comparing higher T_1_ at normoxia than hyperoxia are shown in (d). All maps are shown overlaid on the 1-mm isotropic T_1_-weighted MNI template.

## Discussion

Using an optimised whole-brain SPACE MRI protocol, with a long TE to minimise contributions from brain parenchyma, and a hyperoxia protocol, this study examined how 100% O_2_ inhalation affects the T_1_ of CSF across the whole brain in healthy participants. High-resolution and low-resolution acquisitions, with a long TE, enabled the calculation of 3D T_1_ maps of brain CSF, free from signal contamination from surrounding brain tissue. These maps revealed that O_2_ inhalation causes widespread T_1_ shortening in the cortical subarachnoid space, basilar cisterns, and cisterna magna, but not in all ventricular regions over the 10-15 minute timescale of hyperoxia studied here. Furthermore, in the cortical subarachnoid space and basilar cisterns, O_2_-induced T_1_ shortening of CSF became apparent after approximately 7 minutes of continuous O_2_ breathing and reached significance after approximately 10 minutes (Figures 6d and 6f).

In the present study, the SPACE sequence [24] was specifically optimised to provide 3D acquisitions with sub-millimetric resolution, making it suitable for studying CSF anatomy. Previous studies on hydrocephalus [44,45] have proposed a similar adaptation of the SPACE protocol to illustrate CSF anatomy, but the present study is the first to propose the SPACE protocol for T_1_ mapping in this area. The SPACE sequence was also optimised to achieve a faster acquisition time by lowering the image resolution, enabling the observation of time-dependent CSF T_1_ changes during hyperoxia. The present application of the SPACE sequence represents a flexible tool that can be adapted in future studies, with a focus on the structural aspects of CSF anatomy or the dynamic changes in CSF longitudinal relaxation time constant. While tested on a Siemens system, protocols from other MRI manufacturers offer potential for similar modifications: the volume isotropic turbo spin echo acquisition (VISTA) (Philips Healthcare, Best, The Netherlands) and the CUBE sequence (GE Healthcare, IL, USA).

The SPACE sequence readout presents some limitations as it is inherently affected by both T_1_ and T_2_ decay, with additional contributions from stimulated echoes arising from the reduced flip angle refocussing train [24,46]. In this context, the use of non-180° refocussing pulses causes a portion of the magnetisation to be stored along the longitudinal direction between pulses, where it undergoes T_1_ recovery rather than T_2_ decay [24]. This has two potential consequences for T_1_ mapping: the effective TI experienced by the spins may differ from the nominal TI due to ongoing T_1_ recovery throughout the echo train, and the effective TR may be shortened relative to its nominal value due to the pseudo-steady state reached during the long echo train. Furthermore, the SPACE signal model implemented in our Bloch simulations did not account for additional effects, such as magnetisation transfer or deviations from the nominal refocussing flip-angle train. Therefore, the values reported in the present study should be considered as apparent T_1_ estimates, and future work should aim to account for additional deviations of the SPACE signal evolution from the nominal model. Nonetheless, because the main focus of this work was on assessing T_1_ shortening caused by hyperoxia and T_1_ differences across ROIs, we deemed this apparent T_1_ measure appropriate for the intended objectives. Additionally, the 50-ms T_1_ spacing chosen for the lookup table set the granularity of the simulated TI_2_/TI_1_ ratio and that of the T_1_ curve. Thus, T_1_ values estimated in a single voxel, were quantised at most at 50 ms. For each ROI, the reported T_1_ value was calculated by averaging across 27-29 subjects the median of the T_1_ distribution in the whole ROI. Thus, the 50-ms quantisation effect averaged out, yielding an effective resolution much smaller than 50 ms and well below the group-level differences reported here. Hence, the 50-ms step size for the T_1_ did not limit the detectability of hyperoxia-induced changes.

Having such a protocol readily available on clinical MR systems is timely, given the growing interest in cerebral CSF spaces and CSF circulation [47]. One practical limitation of the proposed protocol is its long duration for high-resolution T_1_ mapping (approximately 15 minutes). However, considering that basic science research is a primary focus of the neurofluids field, we believe that such a long acquisition time in a research setting on healthy subjects is manageable. Future work exploring specific clinical applications of this protocol could refine it for targeted clinical questions within more acceptable examination times.

Regional analysis of T_1_ in CSF-containing structures showed that, at baseline, the median T_1_ of CSF was systematically longer in the anterior lateral ventricles compared to all other regions (Figure 5, normoxia). This result agrees with a previous study showing that CSF in the frontal horns of the lateral ventricles has baseline T_1_ values significantly longer than in the subarachnoid space [13], possibly caused by the lack of blood-CSF water exchange sites in the anterior ventricles. The presence of such sites elsewhere, specifically the choroid plexuses in the ventricles or pial arteries entering the brain from the subarachnoid space, allows for transfer of water from blood, having a relatively short T_1_, to CSF [48]. This mixing effectively shortens the T_1_ of CSF relative to areas without blood-CSF water exchange.

In the cortical subarachnoid space and basilar cisterns, 100% hyperoxia significantly reduced T_1_ compared to baseline values (Figures 5d and 5f) over a timescale of minutes (Figures 6d and 6f). In these ROIs, the observed T_1_ shortening at our chosen timescale was consistent with previous studies applying a 100% hyperoxic challenge for 15 or 30 minutes [15,17]. Additionally, the rapid hyperoxia-induced T_1_ shortening in sulcal CSF was consistent with a study reporting a dynamic signal increase on fast/turbo spin echo images during 5 minutes of 100% hyperoxia [49]. However, previous studies reported no T_1_ change in the lateral ventricles during 100% hyperoxia [15,17,49]. Here, instead, T_1_ was reduced in the posterior lateral ventricles compared to baseline values (Figure 5b), but the timing of this change could not be detected on the lower-resolution data (Figure 6b). Similar discrepancies between significant T_1_ shortening at a higher resolution and no significant T_1_ shortening at a lower resolution were observed in the cisterna magna (Figures 5g and 6g). These results suggest that, in the posterior lateral ventricles and cisterna magna, higher spatial resolution is required to resolve hyperoxia-induced T_1_ changes, likely by reducing partial volume effects from surrounding tissue. An additional factor is that the low-resolution data set lacked a true normoxia baseline (this comparison was only available in four subjects, precluding statistical analysis), meaning that a hyperoxia-induced T_1_ change in these regions may not have been detected. Consistent with the absence of choroid plexuses in the anterior lateral ventricles, no significant T_1_ shortening was observed in this region under hyperoxia, in line with the anatomical explanation for its longer baseline T_1_ (Figures 5a and 6a). In contrast, both the third and fourth ventricles contain choroid plexuses, but neither showed significant T_1_ shortening under hyperoxia: the third ventricle showed no change (consistent with previous results [15]), and the fourth ventricle showed only a non-significant trend in the high-resolution data (Figure 5e; *p*=0.055). This apparent inconsistency with their anatomy likely reflects an insufficient timescale of hyperoxia exposure rather than a true absence of O₂ transfer into these compartments, and is consistent with the general observation that T_1_ shortening in larger or more enclosed ventricular spaces may require longer exposure durations to become detectable. Moreover, differences in observations across studies on the significance of T_1_ shortening in the lateral ventricles at hyperoxia might be due to varying definitions of ROIs within these anatomical regions. One study segmented a region in the anterior horns of the lateral ventricles based on a 20-mm-thick oblique coronal 3D slab [15], while the others segmented a region across the entire lateral ventricles based on a 91.5-mm-thick transverse 3D slab [17] or a single 7-mm-thick transverse 2D slice [49]. In contrast, the present study subdivided the lateral ventricles into anterior and posterior, highlighting that T_1_ shortening related to 100% hyperoxia only occurs over the timescales studied here in the medial-posterior part of the lateral ventricles, where the choroid plexuses are located. The timescale selected in this and earlier studies may have been insufficient to detect T_1_ shortening throughout all large-volume CSF compartments.

During 100% hyperoxia, the larger and faster change in T_1_ in the cortical subarachnoid space and basilar cisterns compared to the posterior ventricular regions (Figures 5b, 5d, 5f, 6b, 6d and 6f) is likely to be related to the nature of blood-CSF exchange sites and the volume of CSF present in each compartment. Notably, even among the ventricular regions, the magnitude and detectability of T_1_ shortening varied: a significant change was detected in the posterior lateral ventricles at high resolution, a non-significant trend was observed in the fourth ventricle, and no change was detected in the anterior lateral ventricles or third ventricle. This gradient is consistent with regional differences in the density and surface area of blood-CSF exchange sites across the ventricular system. These two factors, the nature of the exchange sites and CSF volume, are discussed in more detail below.

Around the cortex, O_2_ from blood enters the CSF via pial arteries on the brain surface, which are part of a network within the subarachnoid space, a CSF-filled space located between the pia mater and arachnoid mater. Pial arteries have thick walls composed of endothelium, basement membranes and layers of smooth muscle [50]. Because, unlike parenchymal arteries and capillaries, pial arteries are directly adjacent to the CSF, a difference of oxygen partial pressure (ΔPO_2_) exists that favours passive outward diffusion of O_2_ into CSF. This diffusion, sometimes termed pre-capillary O_2_ shunting, allows O_2_ to enter the CSF before the blood reaches the capillaries [51]. In the ventricles, O_2_ from blood enters the CSF via the choroid plexus, a highly vascularised structure composed of villi, each containing fenestrated capillaries, stromal tissue, and a layer of choroidal epithelial cells interconnected by tight junctions, which form the BCSFB [52]. O_2_ dissolved in plasma is expected to diffuse rapidly and freely across these structures [52,53].

We suggest that a key factor determining the rate and degree of T_1_ shortening in a given CSF compartment is the relationship between the permeability surface area product of the local BCSFB and the volume of CSF into which O_2_ is transferred. We hypothesise that there is a greater ratio of pial arteries relative to the sulcal CSF volume than there is of choroid plexus vessels relative to ventricular CSF volume, thereby resulting in a faster and greater T_1_ shortening in the cortical subarachnoid space during hyperoxia, consistent with the sulcal-ventricular difference reported previously [17]. In ventricular CSF, the dilution of O_2_, related to the local volume of CSF, is likely to be slower compared to sulcal CSF. This explanation has also been proposed by studies performed in subjects receiving 100% O_2_ under general anaesthesia [54] or through a face mask [55], which showed signal hyperintensities in subarachnoid CSF much higher than in ventricular CSF on fluid-attenuated inversion recovery MRI [54,55]. A similar principle may explain the pronounced T_1_ shortening observed in the basilar cisterns: this compartment surrounds the large-calibre arteries of the circle of Willis and basilar artery at the skull base, which, like pial arteries, are directly adjacent to CSF and offer a substantial surface area for O₂ exchange relative to the local CSF volume. The cisterna magna, in contrast, lacks a substantial local exchange surface of its own, as it contains neither a high density of pial-type arteries nor a choroid plexus. Despite this, it demonstrated a significant, though smaller and slower, T_1_ shortening than the basilar cisterns. The cisterna magna is anatomically continuous with the basilar cisterns, but it also receives a direct influx of unoxygenated CSF from the fourth ventricle via the foramen of Magendie [56,57]. Because the fourth ventricle itself showed no significant T_1_ shortening, it potentially served as a relatively low-O₂ fluid source, being insulated from pial vessels and continuously flushed by fresh upstream CSF. We therefore suggest that the intermediate T_1_ shortening in the cisterna magna reflects a mixing zone, where unoxygenated ventricular outflow blended with heavily oxygenated CSF driven backward from the adjacent basilar cisterns via pulsatile mixing.

In addition to the number of vessels exchanging O_2_, their permeability surface area product will depend on the structure of the vascular walls and thus their permeability. We propose that another factor influencing O_2_ transfer from blood into ventricular CSF is O_2_ consumption within the BCSFB structure. In pial arteries, O_2_ diffuses relatively freely along a steep driving ΔPO_2_. In contrast, at the choroid plexus-CSF interface, epithelial cells are highly active in CSF secretion (via ion pumping) and are rich in mitochondria to support this process [58]. We hypothesise that this high metabolic activity makes the epithelium a metabolic sink, rapidly consuming O_2_ that diffuses into the stroma. Consequently, the final ΔPO_2_ driving O_2_ into the bulk ventricular CSF is reduced due to this consumption, making the transfer rate different from that in the pial artery wall. Furthermore, differences in the baseline PO_2_ within the ventricles and the sulci, driven in part by the continuous circulation and bulk flow of CSF, may also contribute to the local ΔPO_2_ and the effective volume of O_2_ distribution. Consequently, an abnormal hyperoxia-induced T_1_ change in ventricular CSF, larger or smaller than expected based on healthy controls, could reflect altered choroid plexus metabolic activity rather than, or in addition to, changes in vascular permeability. This raises the possibility that ventricular CSF T_1_ under hyperoxia could serve as a non-invasive marker of choroid plexus dysfunction, which has been implicated in several neurological conditions including Alzheimer’s disease, normal pressure hydrocephalus and multiple sclerosis [59].

Based on our observations, we propose that O_2_ delivered via hyperoxia may offer a complementary approach to ASL-based water exchange measurements. In contrast to ASL-based methods, which measure rapid aquaporin-mediated water exchange from blood to CSF on a timescale of seconds, the hyperoxia-induced CSF T_1_ shortening is MR-detectable on a timescale of minutes. We also propose that O_2_ delivered via hyperoxia may be better suited for investigating exchange between blood and CSF compartments compared to GBCAs. Firstly, O_2_ is a simple, rapidly metabolised molecule that avoids the risks of synthetic pharmaceuticals like GBCAs. Secondly, the two compounds differ in their transport mechanisms: O_2_ is a small, uncharged, highly lipophilic gas that rapidly traverses endothelial cell lipid membranes via passive diffusion, reflecting transmembrane permeability. GBCAs, conversely, are significantly larger, polar, hydrophilic molecules with a restricted, slower exchange primarily limited by paracellular transport through tight junctions between endothelial cells [5–7]. Therefore, O_2_ could probe transmembrane exchange, whereas GBCAs probe the size-exclusion properties of tight junctions. Thus, in the future, T_1_ shortening related to the O_2_ transfer rate from pial arteries to CSF (or penetrating arteries and the adjacent perivascular spaces or brain tissue) could serve as a marker of the vessel wall’s effective permeability to O_2_, and by extension, its passive diffusion properties for other small, lipid-soluble molecules. Indeed, we expect the rate of O_2_ loss to be proportional to the permeability-surface area product of the vessel wall, which represents its effective “leakiness” to O_2_. Thus, for a given ΔPO_2_, a higher O_2_ transfer rate implies a higher effective O_2_ permeability. Factors that alter the physical structure of the arterial wall, such as the number of cell layers, cell junction tightness, or the thickness of the barrier, will affect O_2_ movement. Therefore, a change in the O_2_ transfer rate could correlate with a change in the passive permeability coefficient for other small, passively diffusing molecules, potentially establishing O_2_ diffusion as a new marker of vascular structure in pathologies affecting the vasculature, such as dementia [60]. Notably, measurements of the O_2_ transfer rate from pial or penetrating arterioles will only reflect the permeability of the arteriolar wall, which is structurally different from the BBB located at the capillaries. In principle, the rate of O_2_ transfer could be estimated from the temporal evolution of T_1_ shortening by converting the change in longitudinal relaxation between hyperoxia and normoxia to a change in dissolved O_2_ concentration. This conversion is possible using the relaxivity for O_2_ dissolved in CSF (r_1_), that is, the change in longitudinal relaxation rate per unit change in dissolved O_2_ partial pressure, and the solubility coefficient for O_2_ in human plasma at body temperature [61] and dividing by the elapsed time. However, these operations require calibration of r_1_ at 3 T, given that this value depends on the strength of the static magnetic field and has only been calculated at 1.5 T [15] and 7 T [17]. Moreover, they require accounting for concurrent O_2_ consumption by the adjacent brain parenchyma. Therefore, quantifying the absolute transfer rate of O_2_ represents a promising but technically demanding direction for future work.

Potential limitations should be considered when interpreting these results. One factor influencing T_1_ shortening in the cortical subarachnoid space compared to the ventricles could be partial voluming with the surrounding brain parenchyma, which is T_1_-insensitive to hyperoxia [17], and may dilute hyperoxia-induced T_1_ changes in CSF. However, this was addressed by using an extralong TE value to suppress signals from the brain parenchyma around CSF-containing structures (Figure 2), making it unlikely that partial voluming significantly affected the measured T_1_. Furthermore, the influence of hyperoxia on CSF hydrodynamics warrants consideration. Although the effects of normobaric 100% hyperoxia on CSF dynamics in healthy controls have not been investigated, in traumatic brain injury patients, short-term (2 hours) normobaric 100% hyperoxia significantly increased local brain tissue oxygenation without affecting intracranial pressure, cerebral perfusion pressure, or mean arterial pressure [62]. Thus, our sub-30-minute hyperoxia exposure was unlikely to affect CSF dynamics. Additionally, normobaric 100% hyperoxia is known to result in cerebral vasoconstriction, reducing cerebral blood flow by approximately 28-33% in healthy subjects [63]. However, as the hyperoxia-induced T_1_ shortening of CSF is driven by the increased concentration of dissolved O_2_ in CSF (which depends primarily on arterial PO_2_ rather than on blood flow) this effect was unlikely to significantly alter our measurements. Notably, subjects were imaged supine, which enhances CSF flow compared to upright positions [64,65]. Consequently, the observed T_1_ shortening during hyperoxia mainly applies to the supine position used in high-field MRI. Finally, the T_1_ values reported here varied substantially both across CSF regions and across subjects (Figures 5 and 6), likely reflecting a combination of regional and inter-individual physiological differences in CSF properties. The observed inter-regional variability in apparent T_1_ at baseline and the inter-subject variability within individual ROIs both exceeded what could be attributed to differences in dissolved O_2_ concentration alone, given the O_2_ relaxivity values reported for CSF in the literature [15,17] and the arterial PO_2_ increase of up to approximately 400 mmHg expected during 100% O_2_ inhalation relative to room air breathing [17]. This variability means that absolute apparent T_1_ is unlikely to serve as a reliable standalone marker of CSF oxygenation, and the method is better suited to measuring the hyperoxia-induced change in T_1_ relative to a normoxia baseline. This necessarily requires paired acquisitions under both conditions. The high-resolution protocol proposed here resulted in a total scan time of approximately 32 minutes. Although the low-resolution protocol proposed here reduced this time to approximately 7 minutes, it also reduced sensitivity to smaller T_1_ changes, as evidenced by the failure to detect significant shortening in the posterior lateral ventricles and cisterna magna. Additionally, a formal estimate of the minimum T_1_ change detectable at the individual level would require dedicated test-retest data acquired under identical conditions, which were not part of the present study design and represent a direction for future work.

## Conclusions

This study applied a 3D sequence for imaging CSF and measuring CSF T_1_ across the entire brain, adapting a protocol available on clinical MR systems. Experiments conducted with 100% hyperoxia demonstrated that O_2_ induces T_1_ shortening effects in the CSF, these occurring more rapidly in the subarachnoid space and basilar cisterns than in the ventricles. This finding is consistent with the passive diffusion of O_2_ across the high density of blood-CSF exchange sites around the cortex and the skull base. The methodology presented could help characterise CSF structures and assess vascular permeability to O_2_, serving as a marker of cerebrovascular health.

## List of abbreviations

3D: three-dimensional
ANOVA: analysis of variance
ANTs: advanced normalisation tools
ASL: arterial spin labeling
BBB: blood-brain barrier
BCSFB: blood-CSF barrier
CSF: cerebrospinal fluid
FLAIR: fluid-attenuated inversion recovery
FSL: the FMRIB software library
GBCA: gadolinium-based contrast agent
GRAPPA: generalised autocalibrating partial parallel acquisition
IPAD: intramural peri-arterial drainage
LUT: lookup table
MP2RAGE: magnetisation prepared 2 rapid acquisition gradient echoes
MRI: magnetic resonance imaging
O_2_/PO_2_: oxygen/partial pressure of oxygen
ROI: region of interest
SE: standard error
SPACE: sampling perfection with application-optimised contrasts using different flip angle evolution
TE: echo time
TI: inversion time
TR: repetition time
VISTA: volume isotropic turbo spin echo acquisition

## Declarations

### Ethical approval and consent to participate

This study was approved by the Institutional Review Board of the Department of Neurosciences, Imaging and Clinical Sciences, University of Chieti-Pescara (Comitato di Revisione della Ricerca sull’Essere Umano – CReREU, protocol number: 07/2023) on the 16th February 2024. This research was conducted ethically, in accordance with the World Medical Association Declaration of Helsinki. All participants provided written informed consent before enrolment in the study.

### Consent for publication

All participants provided written informed consent to publish the results from this study.

### Availability of data and materials

The anonymised and defaced MR acquisitions supporting the findings from this study are available in this open repository: https://doi.org/10.5281/zenodo.17672756.

### Authors’ contributions

EB was responsible for conceptualisation, data curation, formal analysis, funding acquisition, investigation, methodology, project administration, resources, software, visualisation, and writing (original draft). DDC contributed to data curation, formal analysis, investigation, methodology, software, and writing (reviewing & editing). SP contributed to the investigation and to writing (reviewing & editing). SC contributed to investigation, methodology, and writing (reviewing & editing). EB2 contributed to the investigation and writing (reviewing & editing). MC contributed to writing (reviewing & editing). LC contributed to software and writing (reviewing & editing). GR contributed to software and writing (reviewing & editing). GB contributed to formal analysis, software and writing (reviewing & editing). FG contributed to the investigation and writing (reviewing & editing). ASC contributed to methodology and writing (reviewing & editing). AMC contributed to funding acquisition, methodology, writing (reviewing & editing). RGW contributed to funding acquisition, methodology, project administration, resources, supervision, and writing (reviewing & editing). All authors read and approved the final manuscript.

## Acknowledgements

The authors thank Darien Calvo Garcia and Daniele Petrucci for practical help with the MRI acquisitions, and Domenico Zacà (Siemens Healthcare srl, Milan, Italy) for assistance with the SPACE product sequence.

