## Supplementary Material for "Three-dimensional T_1_ mapping demonstrates the transfer of oxygen into cerebrospinal fluid during hyperoxia" (from another version of this preprint: version 3, 2026-07-06)

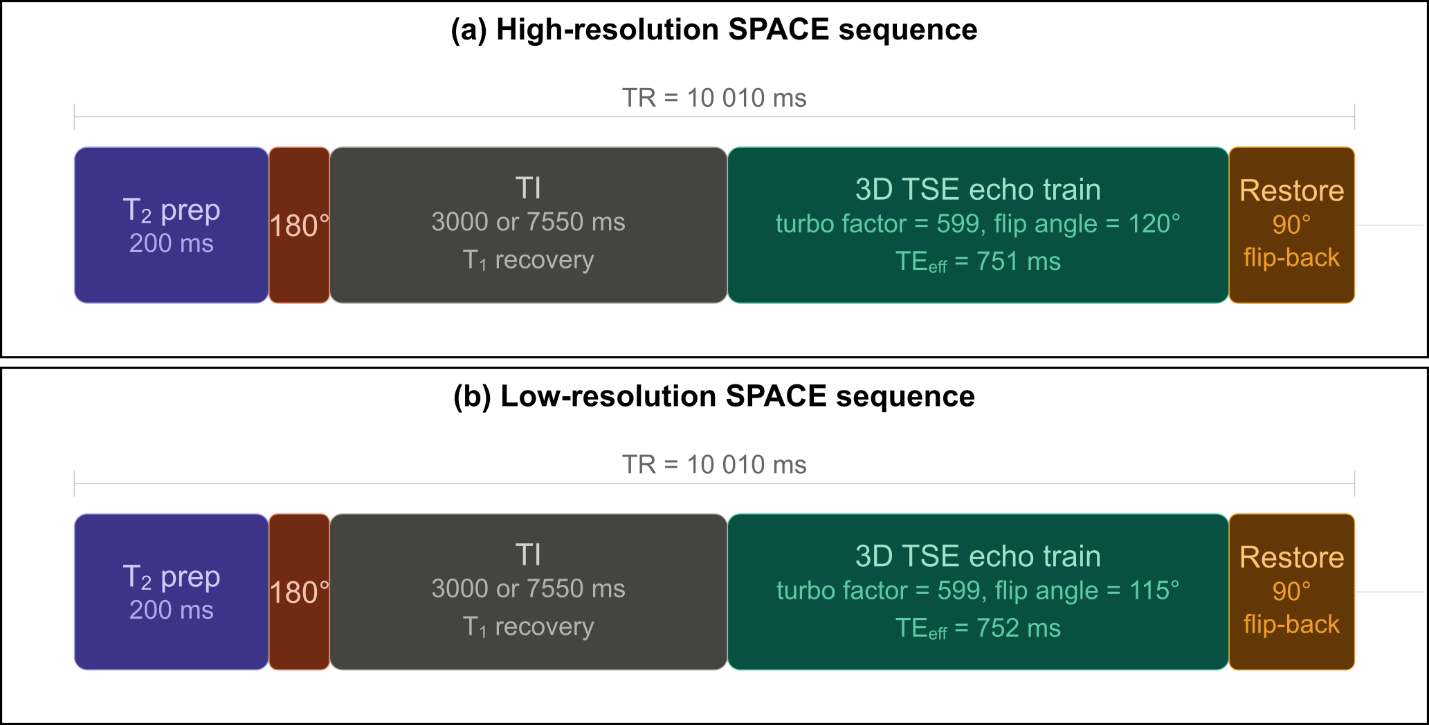

**Supplementary Figure S1. Schematic representation of the SPACE sequence timeline.** The timelines of the two SPACE sequences are shown in (a) for the high-resolution version and (b) for the low-resolution version.

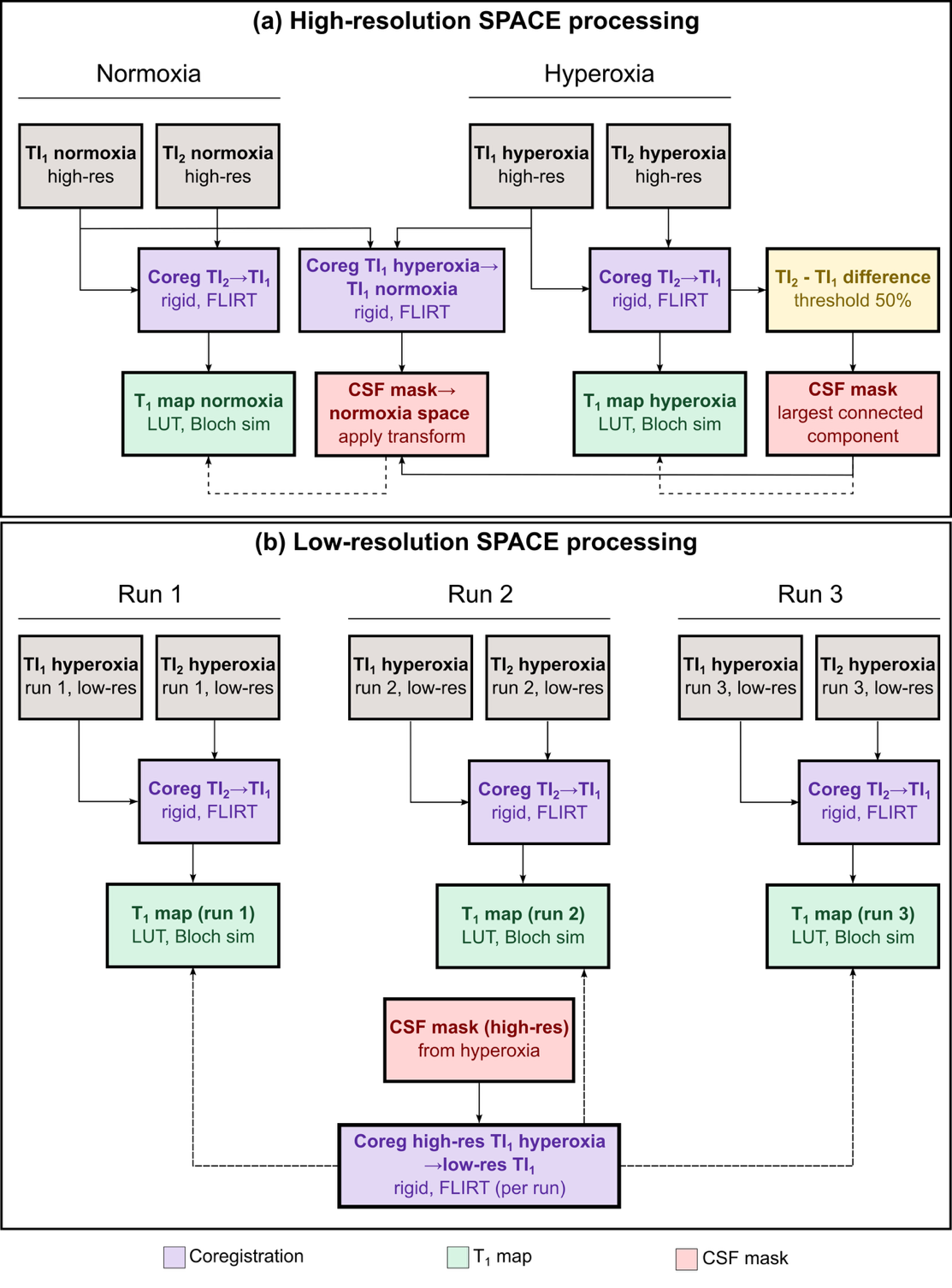

**Supplementary Figure S2. Image analysis pipeline for CSF T_1_ mapping.** (a) High-resolution pipeline: TI₁ and TI₂ images acquired at normoxia and hyperoxia are coregistered, a CSF mask is derived from the hyperoxia difference image, and T_1_ maps are calculated for both conditions using a look-up table from Bloch simulations. (b) Low-resolution pipeline: the same look-up table approach is applied independently to each of the three runs, using a CSF mask propagated from the high-resolution hyperoxia acquisition.

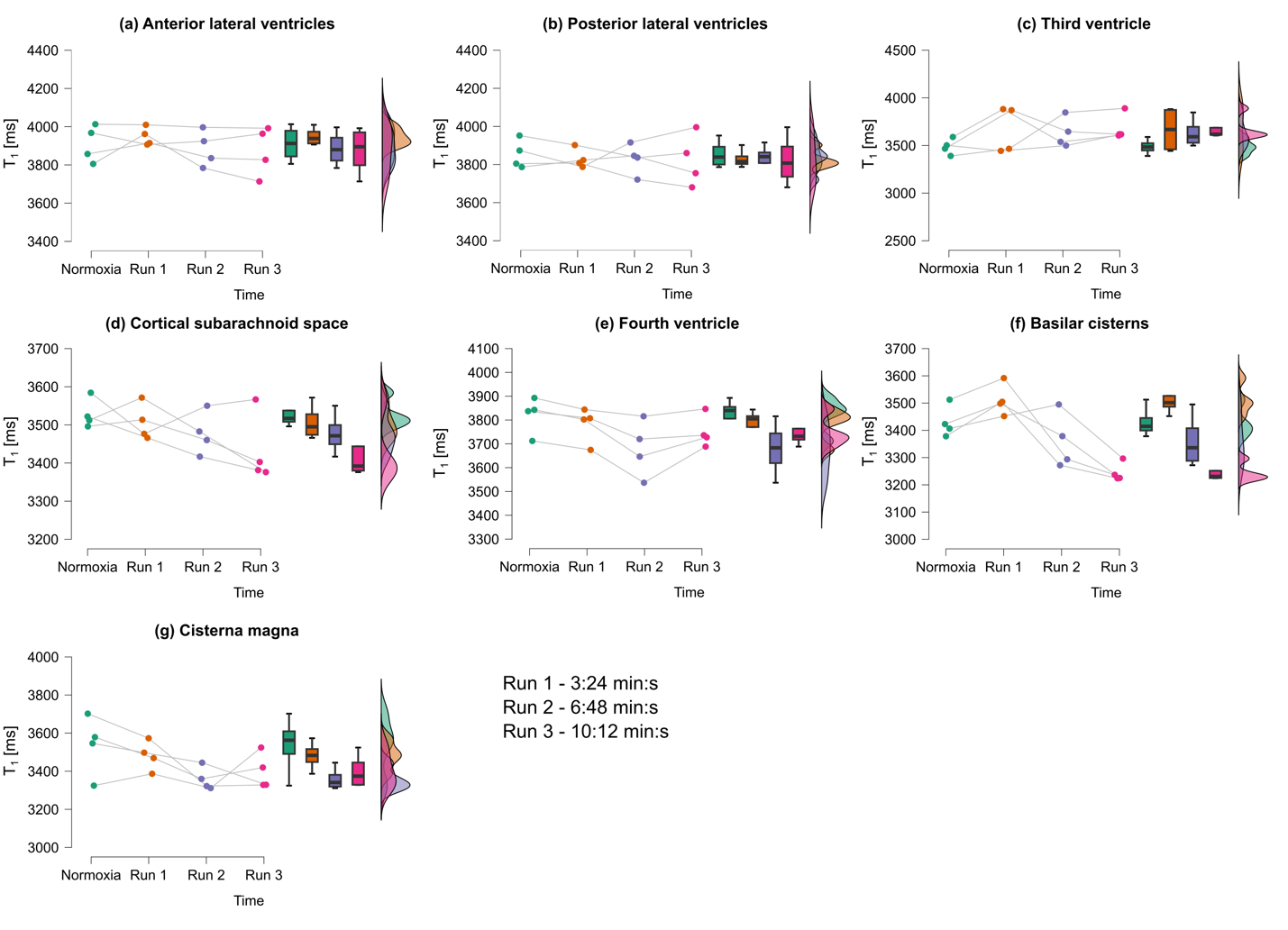

**Supplementary Figure S3. Raincloud plots of regional low-resolution T_1_ measurements including baseline.** For each ROI, the median values of T_1_ measured in four subjects at four time points (normoxia, and runs 1, 2 and 3 during hyperoxia) are shown using a raincloud plot. This plot includes, for each run, a scatter plot of individual data points, a box plot, and a histogram. In the scatter plots for each region, quadruplets of data points connected by a line indicate measurements from the same subject taken at each time point.
